# Mitochondrial genome of *Plasmodium vivax/simium* detected in an endemic region for malaria in the Atlantic Forest of Espírito Santo state, Brazil: do mosquitoes, simians and humans harbor the same parasite?

**DOI:** 10.1101/152926

**Authors:** JC Buery, PT Rodrigues, L Natal, LC Salla, AC Loss, CR Vicente, HR Rezende, AMCR Duarte, B Fux, RS Malafronte, A Falqueto, C. Cerutti

## Abstract

**Background:** The transmission of malaria in the extra-Amazonian regions of Brazil, although interrupted in the 1960s, has persisted to the present time in some areas of dense Atlantic Forest, with reports of cases characterized by particular transmission cycles and clinical presentations. Bromeliad-malaria, as it is named, is particularly frequent in the state of Espírito Santo, with *Plasmodium vivax* being the parasite commonly recognized as the etiologic agent of human infections. With regard to the spatial and temporal distances between cases reported in this region, the transmission cycle does not fit the traditional malaria cycle. The existence of a zoonosis, with infected simians participating in the epidemiology, is therefore hypothesized. In the present study, zoonotic transmission of bromeliad-malaria in Espírito Santo is investigated, based on the complete mitochondrial genome of DNA extracted from isolates of *Plasmodium* species which had infected humans, a simian from the genus *Allouata*, and *Anopheles* mosquitoes. *Plasmodium vivax/simium* was identified in the samples by both nested-PCR and real-time PCR. After amplification, the mitochondrial genome was completely sequenced and compared in a haplotype network, including all sequences of *P. vivax/simium* mitochondrial genomes sampled from humans and simians from all regions in Brazil.

**Results:** The haplotype network demonstrates that humans and simians from the Atlantic Forest share the same haplotype, but some isolates from humans are not identical to the simian isolate. In addition, the plasmodial DNA extracted from mosquitoes revealed sequences different from those obtained from simians, but similar to two isolates from humans.

**Conclusions:** These findings reinforce the hypothesis that in the Atlantic Forest, and especially in the state with the highest frequency of bromeliad-malaria in Brazil, the same parasite species is shared by humans and simians, at least in part. The difference between the sequences of mosquitoes and simians raises two hypotheses: (1) two distinct transmission cycles for human malaria exist in the study area, one of them involving simians and the other exclusive to human hosts, or (2) there is only one transmission cycle involving humans and simians, but the identification of variations among simians was not possible due to a lack of other samples.

## Background

In Brazil, malaria occurs originally across the entire national territory. However, the Amazon region reports 99% of all the cases in the country [1]. Since the 1940s, the national control program has kept malaria transmission restricted to the Northern area. Thus, in the 1960s and 1970s, the extra-Amazonian region came close to complete disease elimination. Nevertheless, residual transmission persisted in areas of the dense Atlantic Forest [2]. In the Atlantic Forest, malaria presents a very low incidence, and the cases are mainly related to *Plasmodium vivax*, presenting few clinical symptoms [1-4]. The low incidence and the territorial dispersion of the reported cases provide evidence in favor of the existence of an unrecognized reservoir of the parasites. This in turn raises hypotheses regarding the participation of asymptomatic carriers or local simians in the transmission [5]. The genetic similarity between the *P. vivax*, that infects humans, and the parasites that infect simians in the Atlantic Forest, named *Plasmodium simium*, reinforces the hypothesis of zoonosis. In fact, most studies indicate that both species are genetically identical [5-8]. However, in a recent study, Brasil *et al.* [9] suggested the possibility of using some single nucleotide polymorphisms (SNPs) in the differentiation of both species.

In the extra-Amazonian region, the term bromeliad-malaria refers to the disease whose vector, recognized as *Anopheles (Kerteszia) cruzii* [10], depends on bromeliads as breeding sites. Molecular and serological evidence presented by different studies has demonstrated that bromeliad-malaria is closely dependent on human activities carried out close to the forest environment [11-13]. In addition, the occurrence of the disease is sparse, and the outbreaks are rare [14]. Considering the characteristics presented above and the fact that the parasites harbored by local simians are genetically indistinguishable from those found in human blood, the hypothesis of a zoonotic scenario for bromeliad-malaria is strongly supported [15-21]. However, even with a variety of scientific investigations corroborating the zoonoses hypothesis, a considerable debate remains regarding the direction of parasite transference. For instance, by comparing the genetic variability in studies based on the Duffy binding protein of erythrocytes collected from simians of the species *Alouatta guariba*, Costa (2014) [22] has suggested that the simian parasite originated from its human counterpart. This hypothesis is additionally supported by Rodrigues *et al.* [23] based on limited genetic variability between *P. simium* and *P. vivax*.

In order to investigate the zoonotic transmission of bromeliad-malaria, this study presents the molecular characterization of *P. vivax/simium* based on the sequencing of the mitochondrial genome of parasites isolated from both human and simian hosts, and, unprecedentedly, from *Anopheles* mosquitoes in an endemic area of the Brazilian Atlantic Forest.

## Methods

### Study area

Espírito Santo is a Brazilian state located in the Southeast region, which harbors large areas of dense Atlantic Forest. The fieldwork for collecting samples of anopheline mosquitoes and monkeys was concentrated in Valsugana Velha, district of Santa Teresa, and the main area with reports of malaria in this municipality. Santa Teresa is located 78 km from the capital of Espírito Santo, Vitória (Figure 1). The landscape in the region is irregular, with a mountainous relief reaching an altitude of 655 meters above sea level, and average temperatures that vary between 15.3 °C and 21.0 °C. Four human blood samples were collected from the inhabitants of Santa Teresa, and 18 from other municipalities of Espírito Santo, also covered by the Atlantic Forest.

**Figure 1.**
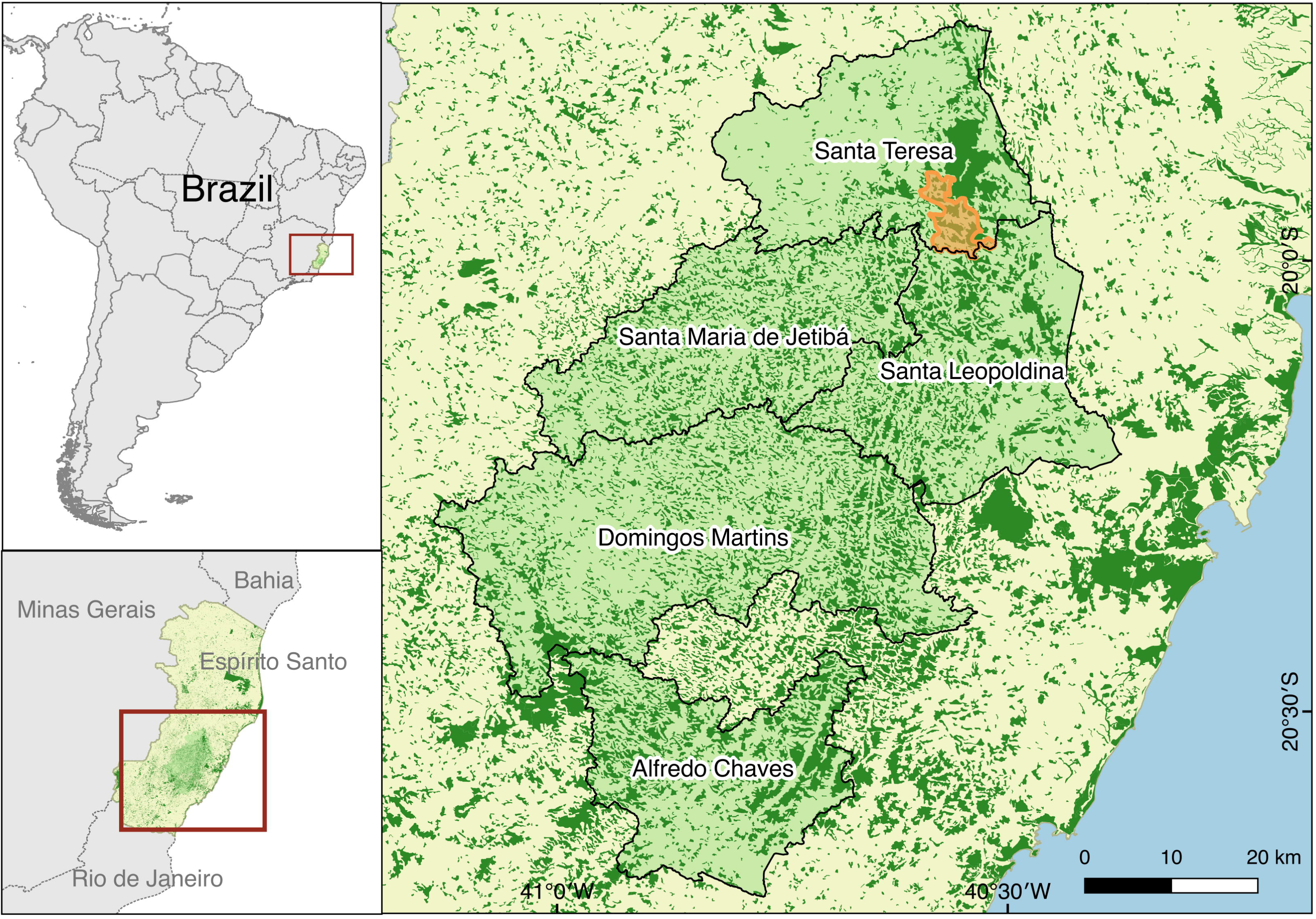
Map showing sampling area of malaria in Santa Teresa municipality, Espírito Santo, Brazil.

### Samples origin

Human blood samples were collected from the 22 cases of malaria caused by *P. vivax* between 2001 and 2004, and previously detected by thick-stained blood smears in the communities of the endemic area [14]. The simian blood sample was obtained from a monkey of the genus *Alouatta* captured alive in Valsugana Velha in 2009. Six specimens of anopheline mosquitoes infected by *P. vivax* and captured in the same area, between 2014 and 2015, were also included (one *Anopheles lutzi*, one *Anopheles strodei* and four *Anopheles cruzii*) [24].

### DNA extraction and confirmation of the infection

Plasmodial DNA from human and simian blood samples was extracted by the QIAamp Blood DNA Mini Kit, while the plasmodial DNA from the mosquitoes was extracted by the DNAeasy Blood and Tissue Kit, both following the instructions of the manufacturer (Qiagen). Infection was confirmed in all samples by nested-PCR [25, 26] and real-time PCR (adaptation from Rubio *et al*. [27]) with primers designed to amplify the 18S RNA subunit gene. Positive and negative controls were used in all reactions.

### Amplification and sequencing of the plasmodial complete mitochondrial genome

The complete mitochondrial genome (6 kb) of *P. vivax*/*simium* from the 22 samples of human blood was amplified and sequenced following the protocol proposed by Rodrigues *et al.* [28]. A new protocol had to be developed in order to perform the amplification of the plasmodial DNA extracted from simian and mosquito samples. Fourteen primers (Pvm1F/Pvm1R to Pvm14F/Pvm14R – Table 1) designed by the software *Primer3* were used in a conventional PCR. The procedure for each sample included 0.5 μl of the enzyme *Taq* DNA polymerase (5.0 U/μl) (Fermentas), 2.0 μl of the extracted DNA, 0.5 μl of each oligonucleotide primer (5.0 μM), 2.0 μl of 10× Buffer for *Taq* DNA polymerase (with KCl), 0.6 μl of dNTP mix (2.0 mM each) and 1.6 μl of MgCl_2_ (25.0 mM), in a final volume of 20 μl. The amplification was run in the GeneAmp PCR 9700 thermocycler (Applied Biosystems), with initial denaturation at 95 °C for 1 minute, followed by 40 cycles of denaturation at 95 °C for 15 seconds, annealing at 60 °C for 30 seconds, and extension for 72 °C for 30 minutes. The final step of the extension was performed at 72 °C for 5 minutes. PCR products were purified by the Illustra GFX PCR and the Gel Band Purification Kit (GE Healthcare Biosciences), and sequenced using the BigDye kit (Applied Biosystems) in the DNA sequencer ABI 3100 (Applied Biosystems). Complete mitochondrial genome assemblies were generated using the software DNASTAR (version 8.1.13, Madison). The sequences were deposited in GenBank (NCBI, https://www.ncbi.nlm.nih.gov/GenBank).

**Table 1.**
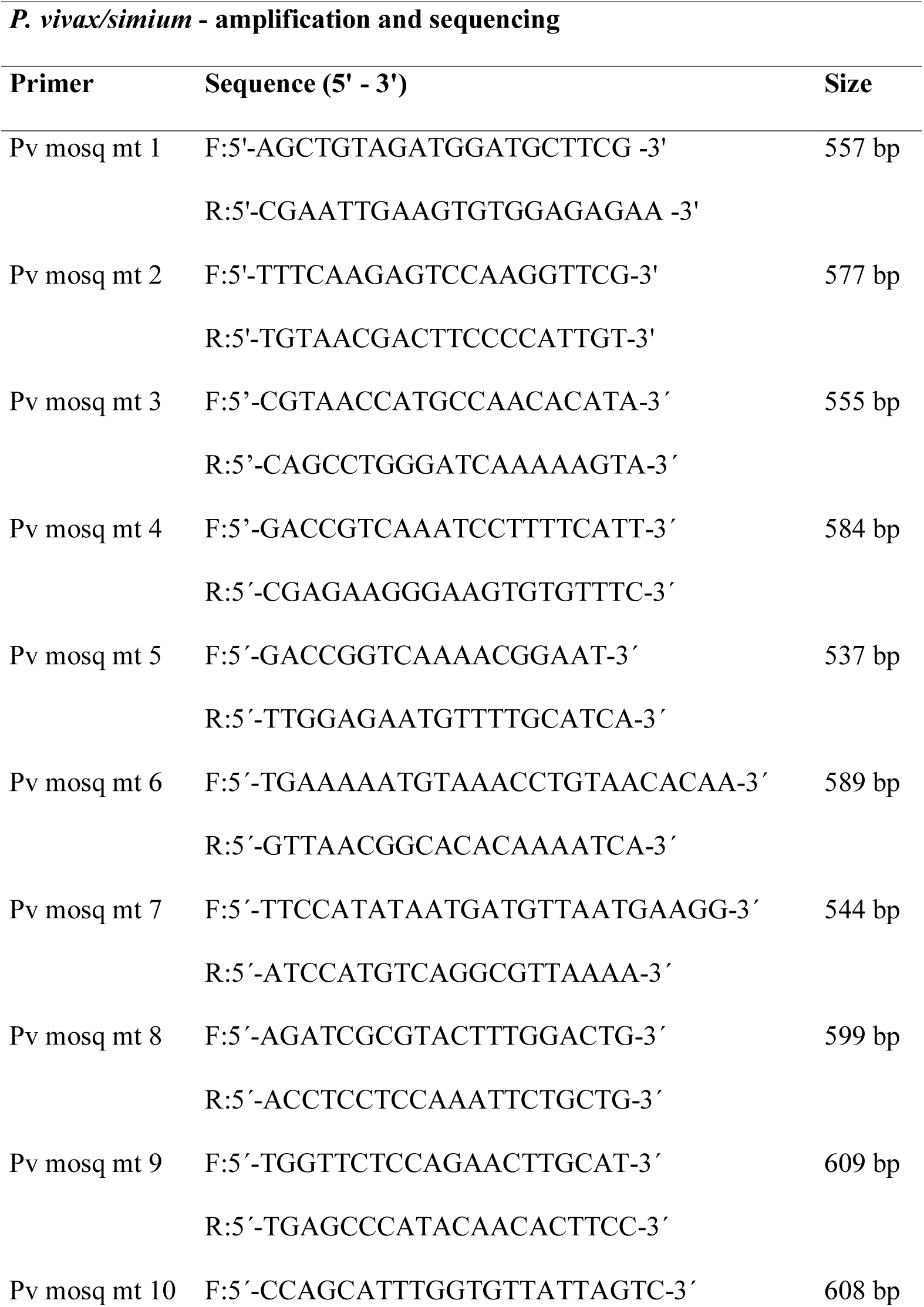
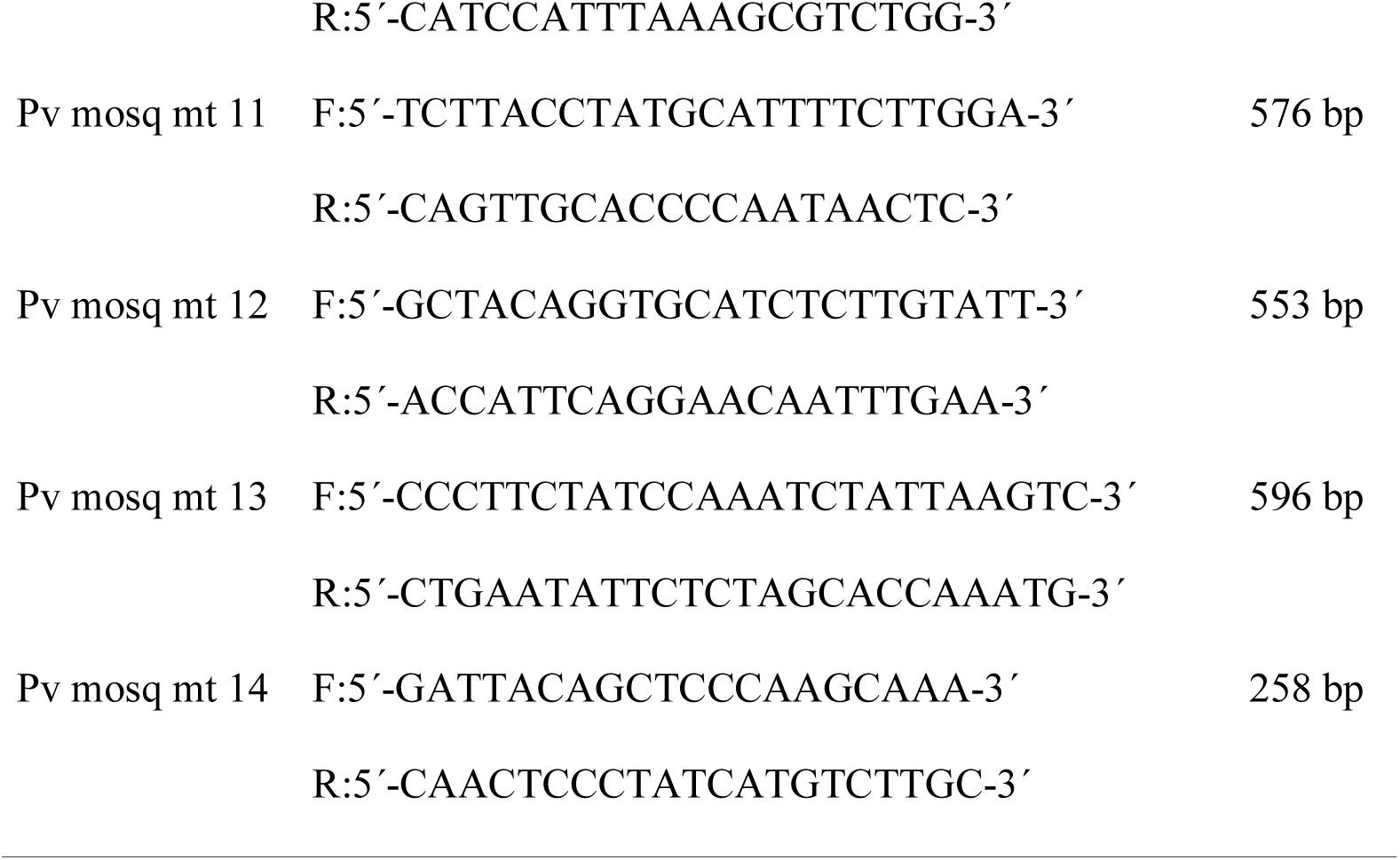
Sequence of primers for amplifying and sequencing of the complete mitochondrial genome of *P. vivax/simium.*

### Data analysis

The sequences of the complete mitochondrial genome were aligned by the program ClustalX (version 2.1) and edited manually in the program MEGA (version 7.0). The genetic p-distance between sequences was also calculated in MEGA. Number of haplotype, haplotype diversity (Hd) and nucleotide diversity (Pi) were calculated using DNAsp (version 5). The haplotype network was generated by median-joining [29] in the program *Network*, version 4.6 (Fluxus Technologies, www.fluxus-engineering.com), with standard parameters. Two different datasets were used for the haplotype network analysis: one including all 29 *P. vivax/simium* mitochondrial genome sequences from Atlantic Forest processed in the present study (n=29); and another also including all 149 *P. vivax/simium* mitochondrial genome sequences from Brazilian Amazonian and extra-Amazonian samples deposited on GenBank, with 10 of the sequences sampled from simian, and 139 from human mitochondrial DNA (mtDNA) (n=179).

Bayesian phylogenetic analysis was carried out for *P. vivax/simium* complete dataset (n=179) using MrBayes (version 3.2.1), with two runs of four chains, three heated and one cold, for 7 · 10^6^ generations. Only groups with Bayesian posterior probability (BPP) ≥ 95% were considered significant. The consensus tree was visualized using FigTree (version 1.4.2).

Taking the GenBank accession number NC_007243.1 as the reference, the SNPs at positions 4134 and 4468 were observed in all sequences sampled from human, simian and mosquito mtDNA, in order to verify if they were useful in distinguishing between *P. simium* and *P. vivax*. As suggested by Brasil *et al.* [9], *P. vivax* should present T/A, and *P. simium* C/G in positions 4134 and 4468, respectively.

## Results

The network comprising only samples from our study (n=29) is shown in Figure 2, and the haplotype network and phylogenetic tree built using the entire database (n=179) are shown in Figure 3. Among the 22 isolates obtained from human blood from Atlantic Forest inhabitants, seven distinct haplotypes were identified (Figures 2 and 3A: Hap1, Hap2, Hap3, Hap4, Hap5, Hap6 and Hap8). Two of them (Figures 2 and 3A: Hap1 and Hap3) were shared with samples isolated from simians. Hap3 has SNPs identical to the *P. simium* sequences deposited on GenBank, as shown in Figure 3A. Hap1 has SNPs identical to the sequence of the isolate obtained from the simian captured in the study area. Two other samples obtained from the human isolates (Figure 3A: Hap8) had SNPs identical to those found in the isolates from human infections acquired in the Amazonian region (*P. vivax*). The remaining four haplotypes (Hap2, Hap4, Hap5 and Hap6) contained SNPs exclusive to the area of the present study (Figure 3A, Table 2).

**Figure 2.**
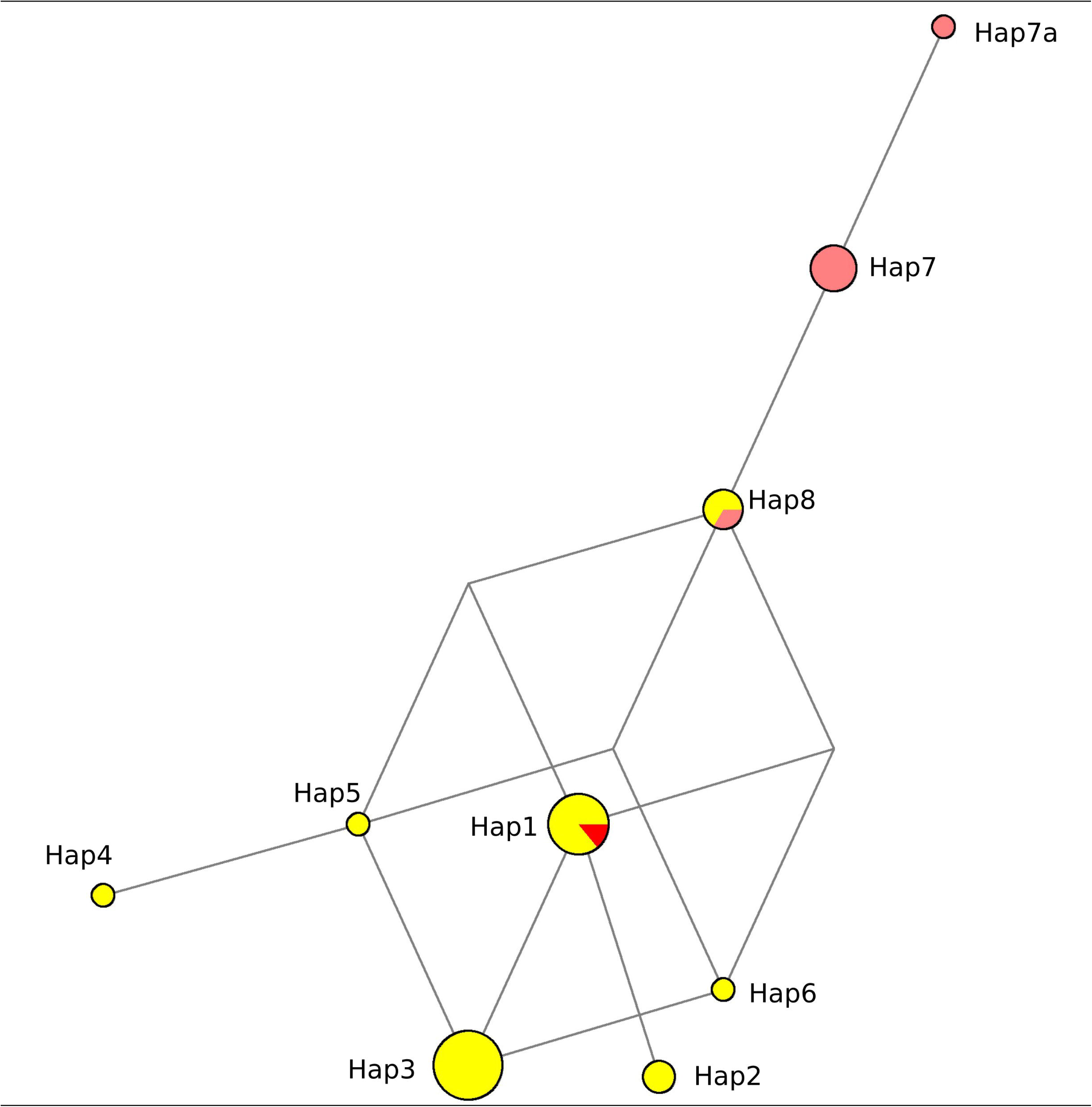
Mitochondrial genome haplotype network of *Plasmodium vivax/simium*, sampled in Atlantic Forest, Espírito Santo, Brazil. Here, 29 samples are presented; 22 from human, 6 from Anopheles mosquitoes and 1 from an *Allouata* simian.

**Table 2.** SNPs of the *Plasmodium vivax/simium* mitochondrial genome from human, simian and mosquito samples from Espírito Santo, Brazil.

|  |  | Source | SNPs (position based on GenBank access NC_007243.1) |  |  |  |  |  |  |
| --- | --- | --- | --- | --- | --- | --- | --- | --- | --- |
| Haplotype | Sample |  | 463 | 1342 | 3325 | 4134* | 4468* | 4511 | 5322 |
| Hap1 | PsimiumES | Monkey | T | C | A | C | G | G | A |
| Hap1 | 1312MT | Human | . | . | . | . | . | . | . |
| Hap1 | 1565MT | Human | . | . | . | . | . | . | . |
| Hap1 | VC57MT | Human | . | . | . | . | . | . | . |
| Hap1 | OJA51_MT | Human | . | . | . | . | . | . | . |
| Hap1 | ACC54_MT | Human | . | . | . | . | . | . | . |
| Hap1 | SV555_MT | Human | . | . | . | . | . | . | . |
| Hap2 | JSB62_MT | Human | . | G | . | . | . | . | . |
| Hap2 | RO54_MT | Human | . | G | . | . | . | . | . |
| Hap3 | GAB847_MT | Human | . | . | T | . | . | . | . |
| Hap3 | 1272MT | Human | . | . | T | . | . | . | . |
| Hap3 | 1411MT | Human | . | . | T | . | . | . | . |
| Hap3 | 1760MT | Human | . | . | T | . | . | . | . |
| Hap3 | 1451MT | Human | . | . | T | . | . | . | . |
| Hap3 | FW63MT | Human | . | . | T | . | . | . | . |
| Hap3 | 143MT | Human | . | . | T | . | . | . | . |
| Hap3 | 111MT | Human | . | . | T | . | . | . | . |
| Hap3 | AJR54_MT | Human | . | . | T | . | . | . | . |
| Hap4 | ALNL53MT | Human | . | . | T | T | . | . | C |
| Hap5 | MA5M61_MT | Human | . | . | T | T | . | . | . |
| Hap6 | 761MT | Human | . | . | T | . | A | . | . |
| Hap7a** | 479mosq | Mosquito | A | . | . | T | A | C | . |
| Hap7 | 485mosq | Mosquito | . | . | . | T | A | C | . |
| Hap7 | 632mosq | Mosquito | . | . | . | T | A | C | . |
| Hap7 | 343mosq | Mosquito | . | . | . | T | A | C | . |
| Hap7 | 260mosq | Mosquito | . | . | . | T | A | C | . |
| Hap8 | 1294mosq | Mosquito | . | . | . | T | A | . | . |
| Hap8 | 40MT | Human | . | . | . | T | A | . | . |
| Hap8 | 103_03MT | Human | . | . | . | T | A | . | . |

\* SNPs suggested by Brasil *et al.* to differentiate between *P. vivax* (T/A) and *P. simium* (C/G).
\*\* Hap7a is represented as Hap7 in the complete data set with 178 samples because position 463 was excluded from the complete database due to missing data in one or more sequences in this site.

As demonstrated in Figure 2, three haplotypes were identified in the isolates obtained from *Anopheles* mosquitoes (Hap7, Hap7a and Hap8): two exclusive to the vector (Figure 2 and 3A: Hap7 and Hap7a), and another (Figure 3A: Hap8) identical to the one identified in the isolates from the Amazonian region, and to two human isolates from the study area (Table 2).

**Figure 3.**
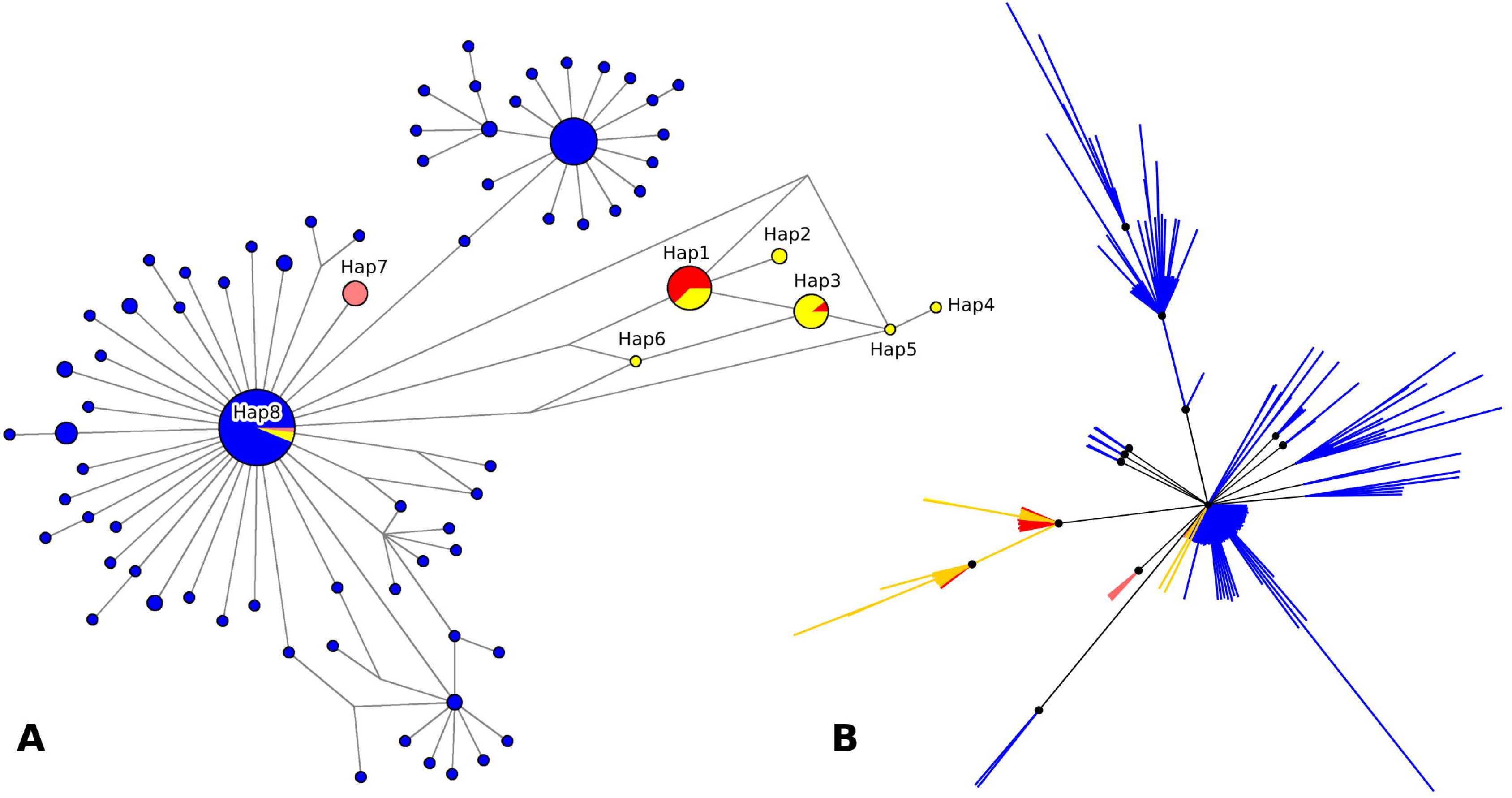
Mitochondrial genome haplotype network and phylogenetic tree of *Plasmodium vivax/simium* from Brazil. 179 samples are presented, including the 29 of Espírito Santo state. (A) The haplotype network by median-joining and (B) the Bayesian phylogenetic tree have the same color pattern, clustered by hosts: blue for human cases from Amazon region; yellow for human cases from Atlantic Forest; red for simian; rose for *Anopheles* mosquitoes. Nodes with Bayesian posterior probabilities ≥ 0.95 are indicated with black circles in the phylogenetic tree.

The genetic divergence within the haplotypes sampled from Atlantic Forest was very low, with only 7 SNPs identified in a total of 5,590 bp (maximum p-distance 0.1%) (Table 2). Among these, 3 SNPs were found in the non-coding region of the mitochondrial genome, 1 SNP within CYTB gene sequence (synonymous mutation) and 3 SNPs within COX1 gene sequence (1 synonymous and 2 nonsynonymous mutations). COX3 sequences were conserved among all samples from Atlantic Forest. Haplotype (Hd) and nucleotide (Pi) diversity were also low for samples from Atlantic Forest (Table 3). Although simian isolates represent larger and more geographically widespread samples than mosquitoes’, haplotype and nucleotide diversity was lower in the former than in the latter (Table 3).

**Table 3.** Number of haplotypes, nucleotide diversity and haplotype diversity in *Plasmodium vivax/simium* mitochondrial genomes sampled from different hosts in Amazonian and Extra-Amazonian regions, Brazil.

| Source of isolates | Number of isolates | Number of haplotypes | Nucleotide diversity (Pi) | Haplotype diversity (Hd) |
| --- | --- | --- | --- | --- |
| Amazonian humans | 136 | 67 | 0.00047 | 0.876 |
| Atlantic Forest humans | 25 | 10 | 0.00031 | 0.823 |
| Mosquitoes | 6 | 3 | 0.00011 | 0.600 |
| Simians | 11 | 2 | 0.00003 | 0.182 |
| Total | 178 | 76 | 0.00051 | 0.907 |

The two SNPs suggested by Brasil *et al.* [9] were not useful to distinguish between *P. simium* and *P. vivax*, at least for the samples of the present study. This is because some haplotypes had sequences different from those proposed as distinctive between *P. simium* and *P. vivax*. More specifically, they showed a combination of these sequences (Table 2: Hap4, Hap5, Hap6). The phylogenetic tree (Figure 3B) also shows that isolates sampled from humans, mosquitoes and simians were not reciprocally monophyletic and some of these sequences clustered together with high statistical support (BPP ≥ 95%).

The haplotype network showed a reticulate relationship between haplotypes, with no evidence of isolation of any haplotype, and with only one or two mutation steps connecting all of the sequences from the samples of the Atlantic Forest region.

## Discussion

*Plasmodium vivax* is a ubiquitous protozoan with a cosmopolitan distribution, causing infections in a number of populations across different continents. Its South American counterpart, *P. simium,* is the etiological agent of malaria in simians inhabiting the Atlantic Forest [30]. Some studies have suggested that *P. vivax* and *P. simium* are the same species, based on their genetic similarities [5-8]. In some areas, including the one of the present study, both simians and humans are infected by this agent, making the hypothesis of zoonosis plausible. The results of the present study, uncovering a haplotype diversity in a situation of low genetic divergence in Espírito Santo, indicate a heterogeneity of the isolates obtained from different host species, and reinforce the understanding that *P. vivax* and *P. simium* are the same species with small genetic variations [31]. These results corroborate the findings of Camargos *et al.* [32], and Rodrigues *et al.* [23], whose phylogenetic analyses of samples from different world regions indicated a recent transfer of the parasite from humans to simians of the New World.

Among the eight haplotypes identified in the study area, two were common to humans and simians, based both on the sequences deposited on GenBank, as well as those obtained from the local simian. This finding represents evidence of parasite transmission from one species to another. At the same time, such a sharing could not be confirmed for the remaining haplotypes, as the sequences were clearly distinctive. Four of the haplotypes obtained from humans were exclusive to the study area, and the two remaining ones were compatible with those previously considered from the Amazonian region [23].

The inclusion of samples obtained from mosquito vectors for the comparison of mitochondrial sequences had never been performed before, despite being previously suggested by Ramasamy [34] and Brasil *et al.* [9]. Interestingly, our results were not consistent with Rodrigues *et al*. [23], who suggested that two SNPs were distinct between malaria from the Atlantic Forest (C/G) and from the Amazonian region (A/T). Rather, plasmodial DNA extracted from mosquito vectors from our Atlantic Forest study area revealed the nucleotides A/T at these loci. These same SNPs were also proposed by Brasil *et al.* [9] as distinctive between *P. vivax* and *P. simium*. However, we demonstrate here that said SNPs were not able to distinguish the two lineages in Espírito Santo, as they were not fixed in at least three samples from humans in the study area (Table 2). Furthermore, the sharing of haplotypes between isolates from different hosts, and the lack of monophyly among samples isolated from humans and simian have been shown in the present work.

The mosquitoes responsible for the transmission of malaria in the Atlantic Forest system belong to the *Kerteszia* subgenus, with the species *Anopheles cruzii* [24, 35, 36] being the most prominent. Specimens of species of the subgenus *Nyssorhynchus* are also captured in the region, occasionally infected by *P. vivax*/*simium* recovered from the blood contained in their abdomens [24]. Despite the possibility of the mosquitoes of the subgenus *Nyssorhynchus* being infected by feeding on human blood, their role as vectors is improbable. Haplotype 8 was obtained from humans and from one pool of *Anopheles (Nyssorhynchus) strodei* captured close to dwellings, suggesting that the mosquitoes were infected accidentally, by the humans. Haplotype 8 has sequences identical to those previously considered specific to the Amazonian region. Two other haplotypes (Hap7 and Hap7a), obtained from other mosquitoes, despite being closely related to those from the Amazonian region, also have distinctive SNPs, making them exclusive for the mosquitoes of the study area (Figure 2, Figure 3, Table 2).

The study has some limitations. The parasite DNA was obtained from humans, the simian, and mosquitoes in different periods, precluding any conclusions regarding a possible circulation of all the haplotypes with the same magnitude at the same time. In addition, only a single simian sample was available, preventing determination of the diversity of the haplotypes infecting this host species (Figure 2). The haplotype network constructed based on 10 simians from the Atlantic Forest revealed two different haplotypes (Figure 3A: Hap1 and Hap3) shared with humans in the study area. Araújo *et al.* [33] highlighted the apparent rareness of simian malaria in the Amazonian region, attributing it to the difficulties in capturing the non-human primates, and in obtaining samples of good quality. The same observational difficulties are applicable to the conditions of the Atlantic Forest.

The presence of sequences identified in isolates obtained from mosquitoes and shared by two isolates from humans, but different from those obtained from simians in Espírito Santo, raises two hypotheses. The first hypothesis is that this is a zoonotic cycle and there are variations in the SNPs among the simians, whose identification was not possible due to the lack of additional simian samples. The second hypothesis is that there are two distinct transmission cycles for human malaria in the study area, one of them involving simians and the other exclusive to human hosts. In order to clarify such questions, future studies should include more samples from simians and vectors, all obtained in the same period.

## Conclusions

Sequencing of the complete mitochondrial genome of *P. vivax/simium* in an area of the Atlantic Forest in Brazil uncovered eight haplotypes, two of which were shared by humans and simians. Interestingly, the other six haplotypes were distinctive, harboring sequences either unique for human infections in the Atlantic Forest or identical to those of the Amazonian region. Such results indicate the possibility of a zoonotic cycle, but due to the differences between plasmodial DNA sequences from simians and mosquitoes, an additional cycle involving only humans cannot be ruled out.

## List of abbreviations

DNA: deoxyribonucleic acid
PCR: polymerase chain reaction
SNP: single nucleotide polymorphism
km: kilometers
°C: degrees Celsius
kb: kilobases
μl: microliters
U: unit
μM: micromolars
mM: millimolars
T: Thymine
A: Adenine
C: Cytosine
G: Guanine
IBAMA: Brazilian Institute of the Environment and Renewable Natural Resources
SISBIO: Biodiversity Information and Authorization System
bp: base pair

## Ethics approval and consent to participate

The collection of human blood samples was performed in a previous study [14] and this material has remained stored since then. In the previous study, blood samples were collected only after obtaining signed informed consent. The collection of samples of simian blood and mosquito specimens [24] had authorization from the Brazilian environmental agency (IBAMA/SISBIO; Number 2508929).

## Consent for publication

Not applicable.

## Availability of data and material

Some datasets analyzed during the current study are not publicly available due to the volume of data, and in deference to other colleagues whose data is not yet published, but are available from the corresponding author upon request.

## Competing interests

The authors declare no competing interests.

## Funding

This research was supported by CNPq/Ministério da Saúde-Decit/ Secretaria de Estado da Saúde do Espírito Santo/ Fundação de Amparo à Pesquisa e Inovação do Estado do Espírito Santo (Grant number 10/2013 PPSUS - 65834119/2014). JCB has a doctoral scholarship from Fundação de Amparo à Pesquisa e Inovação do Estado do Espírito Santo (FAPES, Grant number 139/14). ACL has a postdoctoral scholarship from Fundação de Amparo à Pesquisa e Inovação do Estado do Espírito Santo (CAPES/FAPES, Grant number 68854315/14). The funding body had no influence on the design of the study, the collection, analysis and interpretation of the data, nor on the writing of the manuscript.

## Authors’ contributions

JCB and CCJr conceptualized the main idea of the study. JCB, CCJr and ACL wrote the first version of the manuscript. CCJr, HRR, AMCRD and AF contributed with entomological and epidemiological assistance. JCB, LN and RSM were responsible for the diagnostics of infections. PTR and LCS gave the scientific and technical support for mitochondrial genome amplification and sequencing. JCB, PTH and LCS worked on the laboratory analysis for amplification and sequencing purposes. PTR, ACL and CRV analyzed the sequence data. All authors contributed with a critical reading of the final version of this paper.

## Acknowledgements

We would like to thank the Espírito Santo Health Department for the logistic support in carrying out the fieldwork. Our sincere gratefulness to Dr. Marcelo Urbano Ferreira for the critical reading, donation of reagents and for the always opened-doors laboratory. We especially thank the Santa Teresa population for their warm reception and their trust in our work.

